# The role of *Limch1* alternative splicing in skeletal muscle function

**DOI:** 10.1101/2022.10.05.510856

**Authors:** Matthew S. Penna, George G. Rodney, Rong-Chi Hu, Thomas A. Cooper

## Abstract

Postnatal skeletal muscle development is a highly dynamic period associated with extensive transcriptome remodeling. A significant aspect of postnatal development is widespread alternative splicing changes, required for the adaptation of tissues to adult function. These splicing events have significant implications since the reversion of adult mRNA isoforms to fetal isoforms is observed in forms of muscular dystrophy. LIM and Calponin Homology Domains 1 (LIMCH1) is a stress fiber associated protein that is alternative spliced to generate uLIMCH1, a ubiquitously expressed isoform, and mLIMCH1, a skeletal muscle-specific isoform. mLIMCH1 contains 454 in-frame amino acids which are encoded by six contiguous exons simultaneously included after birth in mouse. The developmental regulation and tissue specificity of this splicing transition is conserved in mice and humans. To determine the physiologically relevant functions of mLIMCH1 and uLIMCH1, CRISPR-Cas9 was used to delete the genomic segment containing the six alternatively spliced exons of LIMCH1 in mice, thereby forcing the constitutive expression of the predominantly fetal isoform, uLIMCH1 in adult skeletal muscle. mLIMCH1 knockout mice had significant grip strength weakness *in vivo* and maximum force generated was decreased *ex vivo*. Calcium handling deficits were observed during myofiber stimulation that could explain the mechanism by which mLIMCH1 knockout leads to muscle weakness. Additionally, *LIMCH1* is mis-spliced in myotonic dystrophy type 1 with the muscle blind-like (MBNL) family of proteins acting as the likely major regulator of *Limch1* alternative splicing in skeletal muscle.

## Introduction

Postnatal skeletal muscle development is a period of dynamic change as the tissue undergoes extensive tissue remodeling to meet new functional demands. A significant aspect of this process is the regulated alternative splicing of hundreds of genes, many of which are temporally coordinated. This process facilitates the transition from fetal to adult protein isoforms integral in the development of adult tissue functionality^1–3^. While large numbers of alternative splicing transitions have been identified in genome-wide transcriptomic analyses, the functional contribution of individual isoform transitions to muscle physiology remains largely unexplored^4–7^. To identify alternative splicing transitions with likely functional importance, we used criteria including conservation of the alternative exons, temporal conservation of splicing pattern transitions, developmental regulation, and coding potential^8–10^. In previous studies, we demonstrated extensive alternative splicing transitions during postnatal skeletal muscle development and redirected splicing of four vesicular trafficking genes toward fetal isoforms in adult skeletal muscle resulted in T-tubule disruption, calcium handling deficits, and impaired force generation^1,11^. Mis-regulated alternative splicing is also known to have pathogenic consequences in skeletal muscle^12^. A notable example is the disease myotonic dystrophy type 1 (DM1), in which perturbed activities of RNA binding proteins Muscleblind-like (MBNL) and CUG-BP Elav-like family (CELF) leads to a reversion to fetal alternative splicing patterns for hundreds of genes, resulting in subsequent muscle wasting^13–15^.

LIM and calponin homology domains 1 (*LIMCH1*) is a ubiquitously expressed gene with mRNA and protein expression in skeletal, cardiac, brain, respiratory, gastrointestinal and reproductive tissues^16,17^. The LIMCH1 protein contains one calponin homology (CH) domain at the N-terminus, several coiled-coil domains located throughout the middle of the protein, and one LIM domain located at the C-terminal region, which canonically have roles in actin binding, protein oligomerization, and protein-protein interactions, respectively^18–20^. The sole study investigating the function of LIMCH1 characterized it as an actin stress fiber associated protein that binds non-muscle myosin 2A (NM2A) activity to regulate focal adhesion formation. Silencing of LIMCH1 led to disrupted focal adhesions and a subsequent increase in cell migration^21^. Several reports have linked LIMCH1 depletion to cancer progression suggesting reduced LIMCH1 levels may serve as a biomarker for tumor growth and/or metastasis in lung, breast and cervical cancers^22–27^. Beyond the role of LIMCH1 in focal adhesion formation and cell migration in vitro, nothing is known about its role in tissue-specific development or physiology.

In adult mouse skeletal muscle, *Limch1* is alternatively spliced leading to the coordinated and skeletal muscle-specific inclusion of six in-frame exons not found in the ubiquitous isoform of LIMCH1, uLIMCH1. This muscle-specific LIMCH1 isoform, termed mLIMCH1, is the predominant isoform in adult skeletal muscle. The six exons introduce 454 amino acids predicted to be disordered with no identifiable domains, which is a common feature of alternatively spliced protein regions^28^. The functional advantage of coordinately including six exons in LIMCH1 to form a novel, skeletal muscle-specific, adult predominant protein isoform is unknown. We hypothesized that mLIMCH1 contributes to adult muscle function and homeostasis based on three key features: developmental regulation involving a fetal to adult isoform transition, highly specific expression of the mLIMCH1 isoform in skeletal muscle, and evolutionary conservation of mLIMCH1 isoform across multiple mammalian species.

In this study, we used CRISPR-Cas9 to delete the genomic segment containing the six skeletal muscle-specific exons of *Limch1* in mice, leading to the constitutive expression of uLIMCH1 throughout postnatal development and in adult mouse skeletal muscle. In mice lacking mLIMCH1, we identified significant muscle weakness *in vivo* without loss of muscle mass as well as significant reductions of maximum force, rate of relaxation, and rate of contraction demonstrated *ex vivo*. We show that both mLIMCH1 and uLIMCH1 preferentially localize to the sarcolemma region in wild-type (WT) myofibers. This localization is disrupted in isolated myofibers in response to the absence of mLIMCH1. Additionally, we found that myofibers lacking mLIMCH1 exhibit disrupted calcium handling, which we propose as a mechanism contributing to muscle weakness. We also show that inclusion of the six skeletal muscle-specific exons requires MBNL activity and LIMCH1 splicing is disrupted in patient-derived DM1 skeletal muscle tissue revealing loss of mLIMCH1 expression as a novel mis-regulated splicing event in DM1 muscle.

## Results

### Alternative splicing generates a skeletal muscle-specific isoform of LIMCH1 during postnatal development

We analyzed our previously published transcriptomic datasets for alternative splicing changes that occur during postnatal skeletal muscle development in mice to identify undescribed protein isoform transitions likely to be important for adult tissue function^1^. We identified a conserved splicing transition in the *Limch1* gene that leads to the coordinated inclusion of six exons that introduces 454 in-frame internal amino acids expressed exclusively in skeletal muscle (**Figure 1A**). The ubiquitous isoform, uLIMCH1, is expressed in most tissues, however in skeletal muscle, mLIMCH1 expression increases soon after birth to become the predominant isoform in adults (**Figure 1B**). The presence and timing of inclusion of the muscle-specific region is conserved between human and mice (data not shown). Reverse transcription (RT)-PCR of WT mouse skeletal muscle tissue indicates a developmental switch from *uLimch1*, the predominant isoform in fetal skeletal muscle, to *mLimch1*, the predominant isoform in adult skeletal muscle (**Figure 1C**). In postnatal day 1 (PN1) skeletal muscle, the six exons unique to *mLimch1* are included in only ∼33% of the total *Limch1* mRNA with *uLimch1* constituting the remaining ∼67% of the *Limch1* mRNA transcripts. However, in adult skeletal muscle, the m*Limch1* isoform constitutes the majority (∼72%) of the total *Limch1* mRNA (**Figure 1C, 1D**).

**Figure 1.**
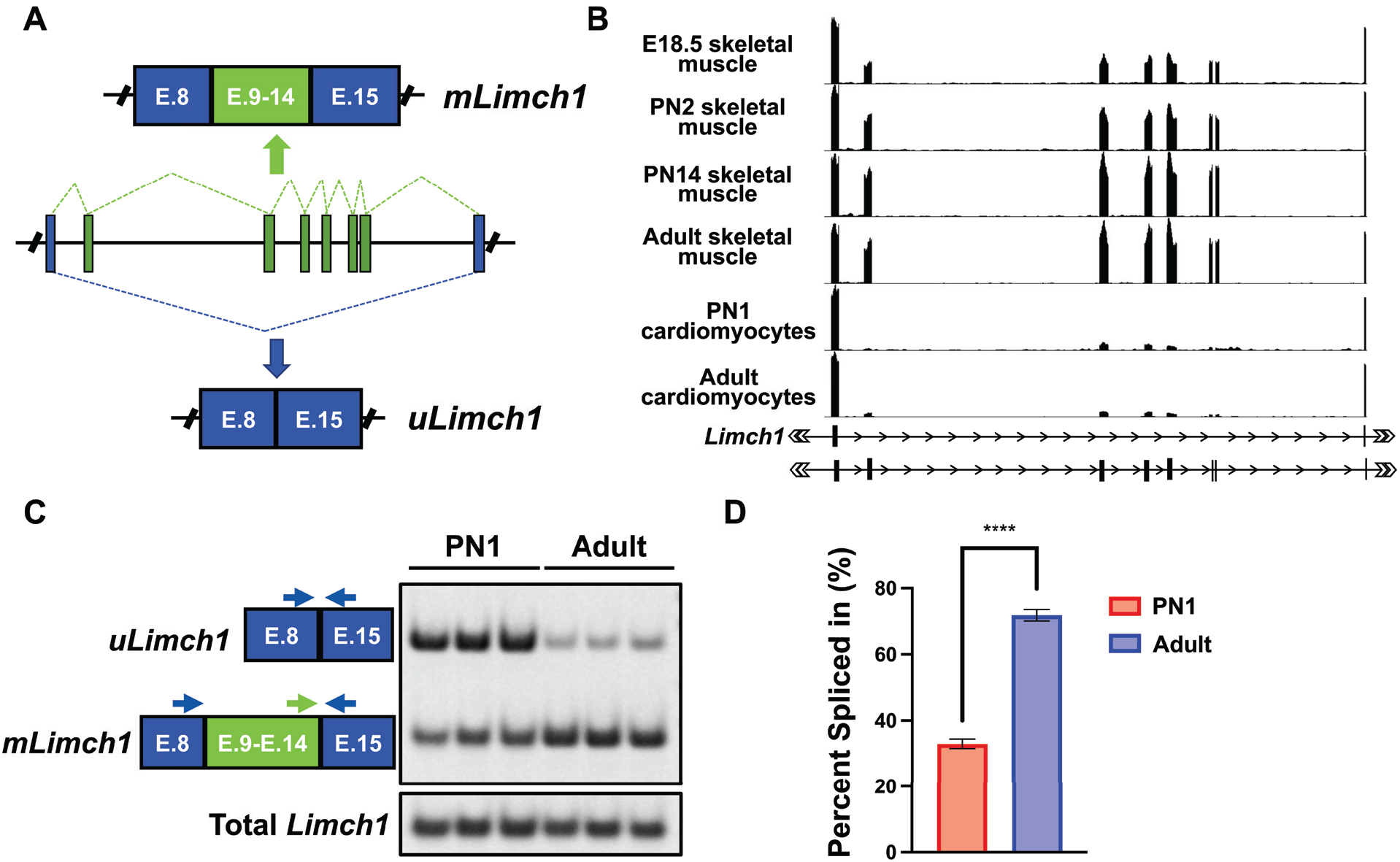
Skeletal muscle-specific inclusion of six exons (E.9-E.14) in *Limch1* during postnatal development. (A) Alternative splicing of the *Limch1* pre-mRNA in skeletal muscle produces a ubiquitously expressed mRNA isoform, u*Limch1* and a skeletal muscle-specific isoform, m*Limch1*. (B) RNA-seq tracks of *Limch1* mRNA from mouse skeletal and cardiac muscle at different ages: E18.5 skeletal muscle, PN2 skeletal muscle, PN14 skeletal muscle, adult skeletal muscle, PN1 cardiomyocytes, and adult cardiomyocytes^1,2^. (C) RT-PCR showing the *Limch1* alternative splicing pattern in PN1 and adult skeletal muscle tissue. (D) Quantification of the *Limch1* muscle-specific exon inclusion in PN1 and adult skeletal muscle (n= 3 PN1, 3 adult). Data represents the mean ± SEM and were analyzed using a two-tailed *t* test. ****P<.0001. E, exon; PN1, postnatal day 1; PN2, postnatal day 2; E 18.5, Embryonic day 18.5; PN14, Postnatal day 14.

### CRISPR mediated deletion of the skeletal muscle-specific exons in *Limch1*

To determine the extent to which the developmentally regulated and conserved splicing of *Limch1* contributes to skeletal muscle physiology, we used CRISPR/Cas9 to delete *Limch1* exons 9-14, that encode for the *mLimch1* isoform from the genome of FVB mice, thereby leading to the expression of only *uLimch1* in adult skeletal muscle of mice homozygous (HOM) for the deleted allele (*Limch1* 6exKO) (**Figure 2A**). RNA-sequencing data confirmed that the six exons unique to *mLimch1* are not included in *Limch1* mRNA from HOM *Limch1* 6exKO mice (**Figure 2B**). Using an antibody that recognizes both isoforms, we performed western blotting for endogenous uLIMCH1 (∼135 kDa) and mLIMCH1 (∼200 kDa) protein levels in different skeletal muscle tissues (**Figure 2C**). HOM *Limch1* 6exKO mice lack expression of mLIMCH1 and exclusively express uLIMCH1. As expected from mRNA data, mLIMCH1 protein is undetectable in WT cardiac and brain tissue (**Figure 2D**). Two independent founder lines from the CRISPR/Cas9 deletion of *Limch1* exons 9-14 were generated and shown to have comparable phenotypes, ruling out the likelihood of CRISPR/Cas9 mediated off-target effects. Additionally, the line of *Limch1* 6exKO mice used for experiments were derived from F1 animals then backcrossed for seven generations to out-cross off-target loci. One of the lines was selected for the subsequent phenotyping assays.

**Figure 2.**
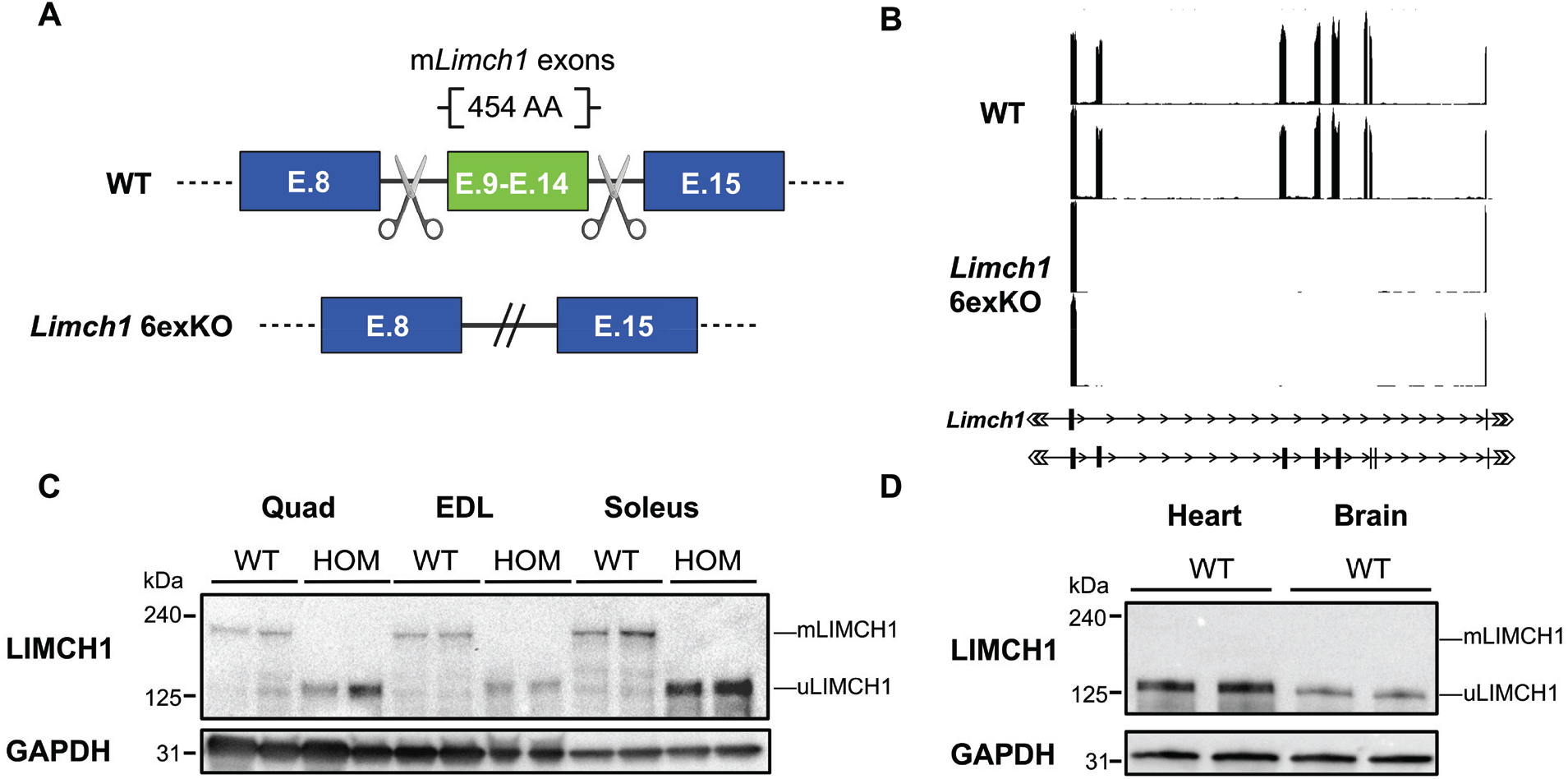
CRISPR/Cas9-mediated germline deletion of exons 9-14 to force the expression of the fetal predominant isoform, uLIMCH1. (A) Diagram of the CRISPR/Cas9 approach to delete *Limch1* muscle-specific exons 9-14 from the mouse genome. Scissors represent the guide RNAs targeting the intronic regions flanking the *mLimch1-*specific exons. (B) RNA-seq tracks of *Limch1* in WT and HOM *Limch1* 6exKO skeletal muscle highlighting the removal of the six skeletal muscle-specific exons in *Limch1*. (C) LIMCH1 protein isoform expression in adult HOM *Limch1* 6exKO mice and WT age-matched controls in quad, EDL, and soleus skeletal muscle (n= 2 WT, 2 HOM). (D) Western blot showing LIMCH1 protein expression patterns in the heart and brain of WT mice (n= 2 WT). E, exon; AA, amino acid; HOM, homozygous; WT, wild-type; quad, quadriceps; EDL, extensor digitorum longus.

### Exclusive expression of uLIMCH1 in adult skeletal muscle leads to reduced grip strength and muscle force generation

To assess the physiological consequences of the loss of mLIMCH1 in adult skeletal muscle, we evaluated muscle function of HOM *Limch1* 6exKO mice side-by-side with WT age-matched controls. We first measured grip strength in adult HOM *Limch1* 6exKO mice. Results showed a significant decrease in the grip strength of HOM *Limch1* 6exKO mice with a decrease of 31% and 29% in male and female mice, respectively (**Figure 3A**). We evaluated maximum force production in both the extensor digitorum longus (EDL) and soleus of HOM *Limch1* 6exKO mice and found a statistically significant decrease in the maximum force produced with a particularly stronger decrease in force generation at higher frequencies (**Figure 3B, 3C**). We next conducted isometric *ex vivo* force analysis of contractile parameters on soleus (predominantly slow-twitch fibers) and EDL (primarily fast-twitch fibers) muscle to directly measure multiple components of muscle function. The rate of relaxation and rate of contraction of a 1 Hz stimulus leading to a twitch was significantly reduced in the EDL from HOM *Limch1* 6exKO mice compared to WT age-matched controls (**Figure 3D, 3E**). The rate of contraction and rate of relaxation in the soleus was not significantly different between HOM *Limch1* 6exKO and WT mice (**Figure S1A, S1B**). No differences in fatigue or recovery after repeated stimulation were observed in EDL or soleus tissue from HOM *Limch1* 6exKO mice (**Figure S1C-S1H)**.

**Figure 3.**
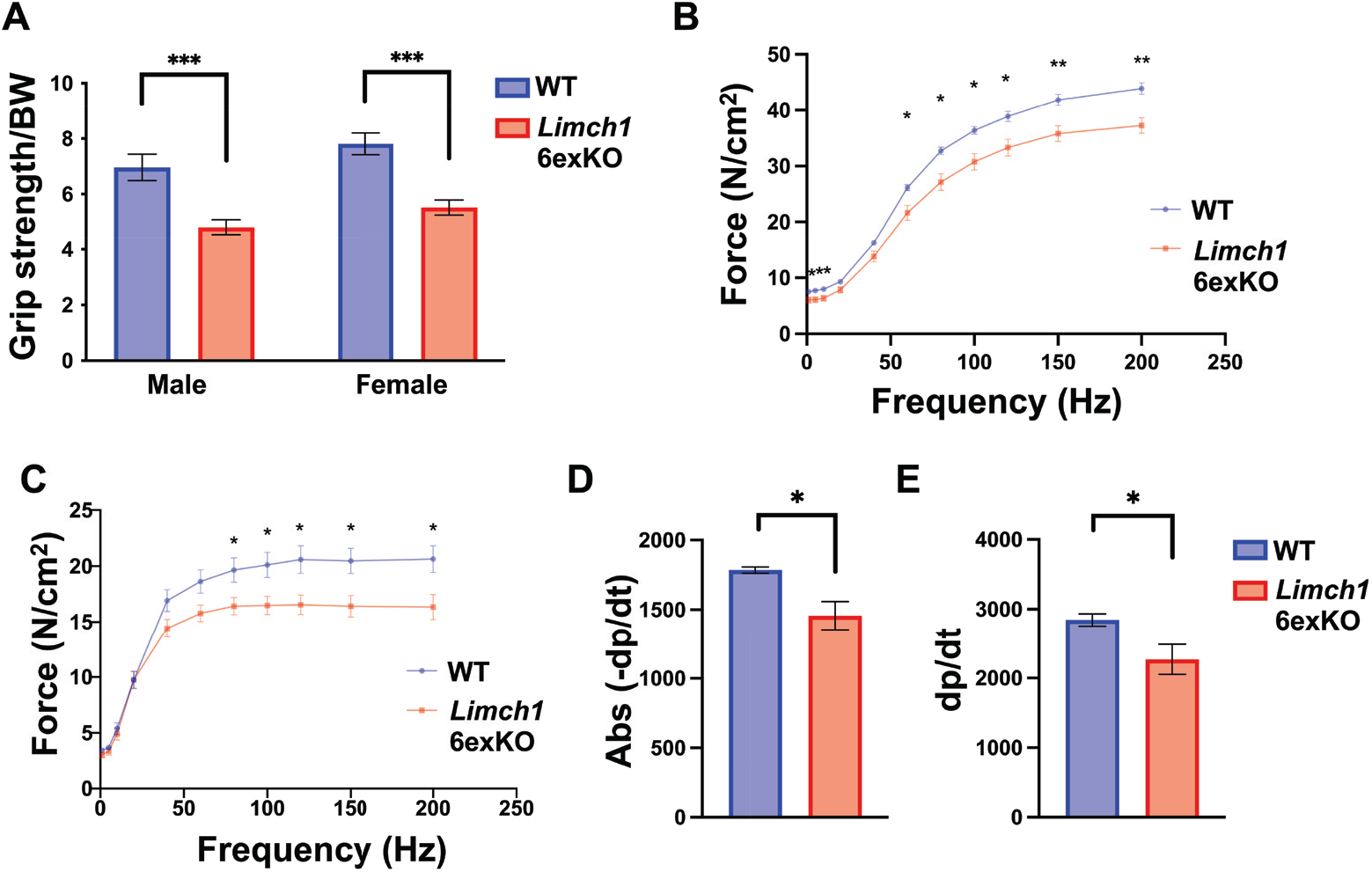
Loss of the muscle-specific LIMCH1 isoform produces grip strength weakness and force production deficits in HOM *Limch1* 6exKO mice. (A) Four-limb grip strength assessment in HOM *Limch1* 6exKO mice and WT age-matched controls (n= 13 WT, 13 HOM males; 12 WT, 14 HOM females). (B) Rate of relaxation (-dp/dt) in the EDL following an *ex vivo* isometric twitch. (C) Rate of contraction (dp/dt) in the EDL following an *ex vivo* isometric twitch. (D) Force-frequency curve during isometric stimulation of the soleus stimulated at increasing frequencies. (E) Force-frequency curve during isometric stimulation of the EDL stimulated at increasing frequencies (For B-E, EDL: n= 6 WT, 8 HOM; soleus: n= 6 WT, 6 HOM). Data represents the mean ± SEM and were analyzed using a two-tailed *t* test. Force frequency curves were analyzed using multiple *t* tests for each frequency tested. *P<.05, **P<.01, ***P<.001. BW, body weight; Abs, absolute value; WT, wild-type; HOM, homozygous; EDL, extensor digitorum longus.

We conducted H&E and picrosirius red staining on muscle cross-sections collected from HOM *Limch1* 6exKO tissue to determine whether underlying histological defects could be contributing to the defects in muscle function. HOM *Limch1* 6exKO mice did not exhibit any differences in myofiber diameter, centralized nuclei, or fibrotic scarring in comparison to WT control mice. Additionally, no differences in muscle weights were observed between the HOM *Limch1* 6exKO and WT control mice. (**Figure S.2**). These results indicate that muscle weakness did not correlate with skeletal muscle histopathology or muscle loss suggesting an intrinsic defect in muscle contraction.

### Localization of LIMCH1 is disrupted in HOM *Limch1* 6exKO myofibers

To elucidate whether loss of the muscle-specific exons impacted LIMCH1 localization we isolated flexor digitorum brevis (FDB) myofibers from WT and HOM *Limch1* 6exKO mice and performed immunofluorescence staining to identify endogenous localization patterns of mLIMCH1 and uLIMCH1 using an antibody that recognizes a shared epitope. The strongest LIMCH1 signal in WT myofibers is observed at the sarcolemma region. In HOM *Limch1* 6exKO myofibers, LIMCH1 staining is observed at the sarcolemma region as well but displays increased distribution throughout the myofiber compared to WT myofibers (**Figure 4A**). We quantified the immunofluorescence signal with plot profiles collected along the sarcolemma of the myofiber (along red arrows in **Figure 4A**) which showed that LIMCH1 is distributed in a punctate manner with maximum peak intensity occurring every two microns in both WT and HOM *Limch1* 6exKO myofibers (**Figure 4B**, left panels). We also conducted plot profiles to compare sarcolemmas in WT and HOM *Limch1* 6exKO myofibers (along red line in **Figure 4A**). The signal intensity in WT myofibers was stronger along the sarcolemma compared to the cytoplasm of the myofiber whereas HOM *Limch1* 6exKO myofibers had a dampened signal intensity along the sarcolemma with a more even signal distribution throughout the myofiber (**Figure 4B**, right panels). We analyzed LIMCH1 immunofluorescence throughout the entire myofiber to determine whether the LIMCH1 signal was distributed differently in WT and HOM *Limch1* 6exKO myofibers as the plot profile comparing sarcolemmas suggested. After determining the signal area by thresholding the image and projecting the outline back onto the original image for analysis, we calculated the standard deviation of each pixel (signal) within the myofiber to quantitate signal distribution (**Figure 4C**). The standard deviation of the signal from WT myofibers was increased significantly compared to HOM *Limch1* 6exKO myofibers indicating that in WT myofibers, the protein localization is clustered closer together. Conversely, the lower standard deviation in HOM *Limch1* 6exKO myofibers indicates a broader distribution of LIMCH1 throughout the myofiber and therefore less clustering of LIMCH1 near the sarcolemma (**Figure 4D**).

**Figure 4.**
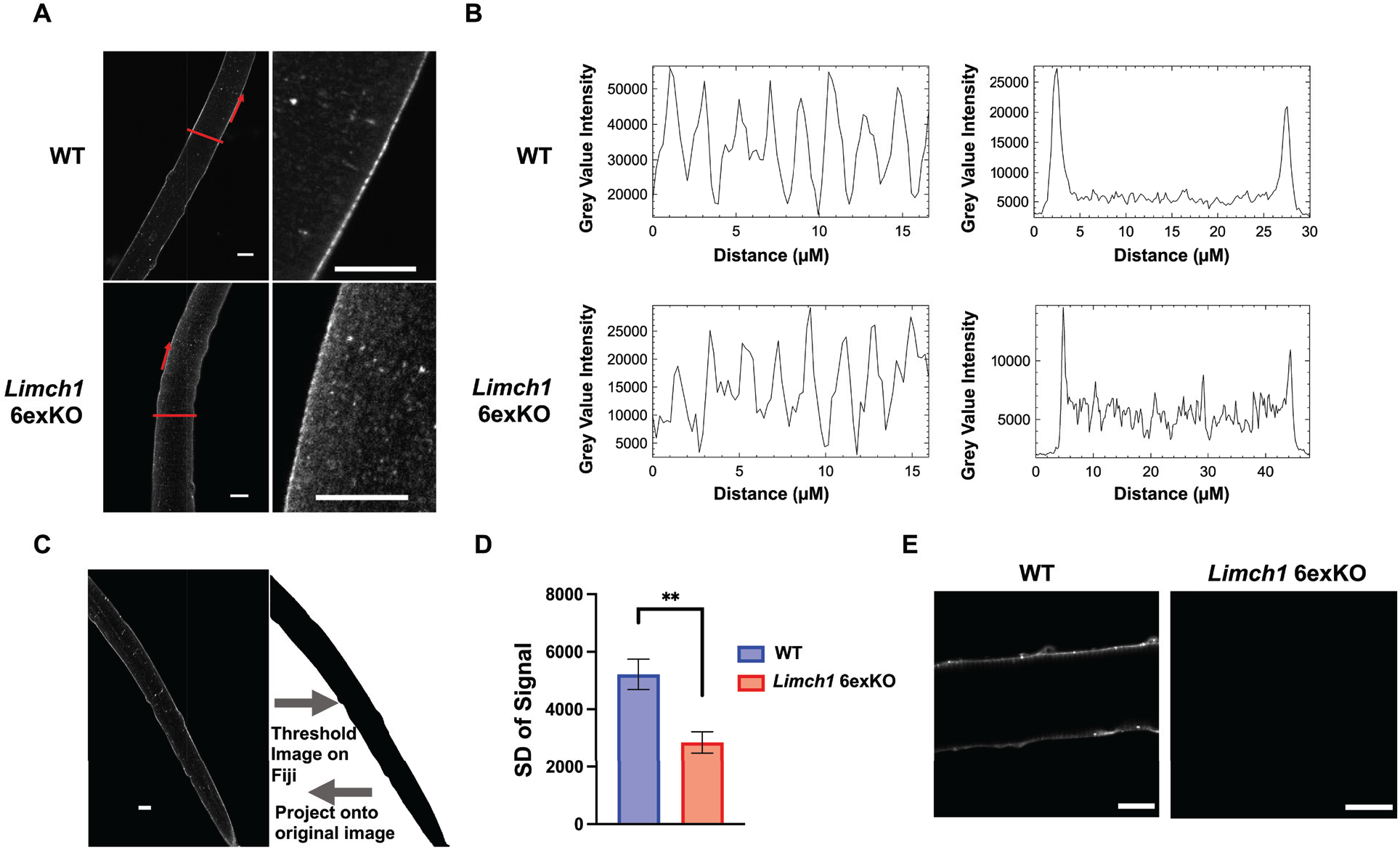
LIMCH1 is mis-localized in HOM *Limch1* 6exKO myofibers. (A) Immunofluorescence staining of endogenous LIMCH1 in WT and HOM *Limch1* 6exKO FDB myofibers with zoomed in panels on the right. Scale bar, 15 μM. (B) Representative signal intensity plot profile along the sarcolemma of WT and HOM *Limch1* 6exKO myofibers (red arrows in A, left panel of B) and from sarcolemma to sarcolemma of the myofiber in WT and HOM *Limch1* 6exKO myofibers (red line in A, right panel of B). (C) Approach for analyzing signal distribution using Fiji to turn the image into a binary image and project the identified signal onto the original image for analyses. Scale bar, 15 μM. (D) Standard deviation of overall LIMCH1 signal in WT and HOM *Limch1* 6exKO myofibers from isolated FDB myofibers (myofibers averaged from n=6 WT, 6 HOM *Limch1* 6exKO). (E) Immunofluorescence staining of endogenous mLIMCH1 in WT and HOM *Limch1* 6exKO FDB myofibers. Scale bar, 15 μM. Data represents the mean ± SEM and was analyzed using a two-tailed *t* test. **P<.01. WT, wild-type; HOM, homozygous; SD, standard deviation; FDB, flexor digitorum brevis.

We developed an mLIMCH1-specific antibody to differentiate uLIMCH1 and mLIMCH1 isoforms and better understand the precise localization of mLIMCH1 in skeletal muscle myofibers. The antibody was commercially synthesized and we subsequently validated the antibody for western blot and immunofluorescence (**Figure S3**). Myofibers stained with the mLIMCH1-specific antibody revealed that mLIMCH1 localization heavily favors the sarcolemma region indicating that uLIMCH1 is unable to recapitulate normal LIMCH1 localization upon mLIMCH1 knockout (**Figure 4E**). Taken together, these results demonstrate that mLIMCH1 preferentially localizes to the sarcolemma region of skeletal muscle myofibers with subsequent mis-localization of the fetal predominant isoform in HOM *Limch1* 6exKO myofibers.

### Intracellular calcium handling is altered in HOM *Limch1* 6exKO skeletal muscle

We conducted *ex vivo* calcium analysis during a myofiber twitch and tetanus to investigate potential contributions of altered calcium handling to the skeletal muscle dysfunction observed *in vivo* and *ex vivo*. Calcium directly regulates muscle contraction while defects in calcium handling can impact force generation and lead to muscle weakness^29–31^. We found that peak calcium levels were decreased in male and female HOM *Limch1* 6exKO derived myofibers stimulated at 1 Hz (**Figure 5A**), 20 Hz (**Figure 5B**), and 100 Hz (**Figure 5C**) frequencies compared to WT myofibers. Results showed an average peak calcium decrease of 26.1%, 23.8%, and 16.0% in male derived myofibers at 1 Hz, 20 Hz, and 100 Hz, respectively, while results showed an average peak calcium decrease of 24.4%, 16.1%, and 13.2% in female derived myofibers at 1 Hz, 20 Hz, and 100 Hz, respectively. We followed up the peak calcium analysis with immunofluorescence of different T-tubule related proteins to determine whether T-tubule structure is intact in HOM *Limch1* 6exKO myofibers and found intact T-tubule structure (**Figure S4**). We find that in the absence of mLIMCH1, calcium handling is disrupted in skeletal muscle while the structural integrity of the T-tubule system remains intact.

**Figure 5.**
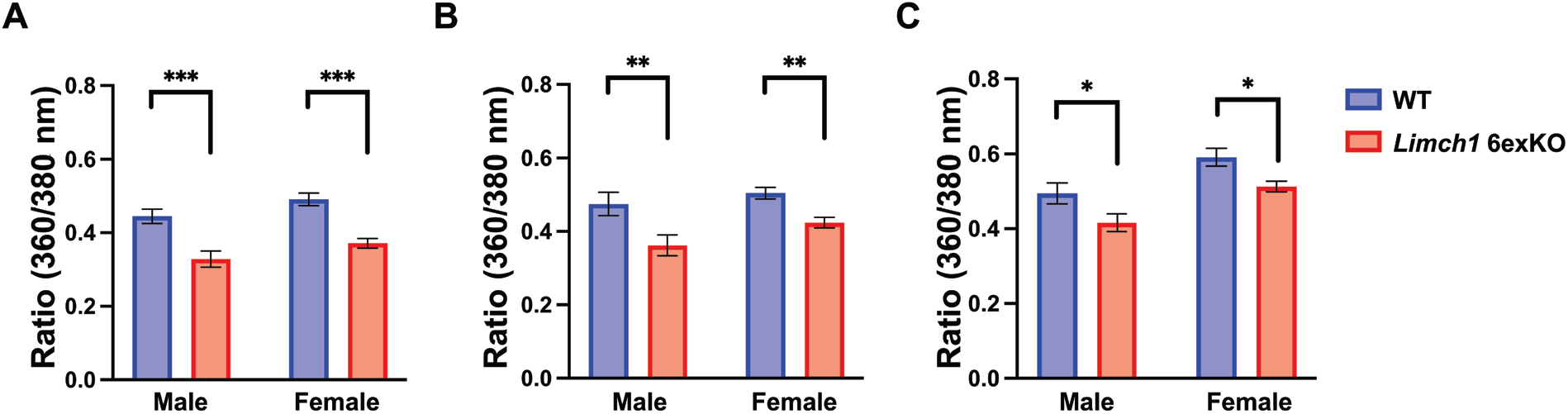
Calcium handling is disrupted in HOM *Limch1* 6exKO isolated myofibers. (A) Peak Fura ratio (360/380 nm) in FDB myofibers stimulated with a 1 Hz twitch. (B) Peak Fura ratio (360/380 nm) in FDB myofibers stimulated with a 20 Hz tetanus. (C) Peak Fura ratio (360/380 nm) in FDB myofibers stimulated with a 100 Hz tetanus (2-3 fibers averaged from n= 10 WT, 11 HOM *Limch1* 6exKO males; 9 WT, 9 *Limch1* 6exKO females). Data represents the mean ± SEM and were analyzed using a two-tailed *t* test. *P<.05, **P<.01, ***P<.001. nm, nanometer; WT, wild-type; FDB, flexor digitorum brevis; HOM, homozygous.

### *LIMCH1* is aberrantly spliced in DM1

Aberrant alternative splicing is a hallmark of many skeletal muscle diseases, with many of the affected genes expressing specific isoforms tailored for skeletal muscle function^13,32,33^. We found that the *LIMCH1* pre-mRNA is mis-spliced in skeletal muscle tissue from adults affected with DM1. DM1 skeletal muscle lacks the appropriate inclusion of the *LIMCH1* skeletal muscle-specific exons and expresses the fetal isoform, *uLIMCH1*, as the predominant isoform (**Figure 6A**). MBNL1 and MBNL2 are RNA binding proteins that regulate hundreds of genes through alternative splicing and MBNL protein sequestration is directly involved in DM1 pathogenesis^34,35^. To better understand the mechanism by which *LIMCH1* alternative splicing is regulated, we looked at LIMCH1 expression in floxed (fl) Ckm Cre; *Mbnl1*^fl/fl^, *Mbnl2*^fl/fl^ mice (**Figure S5**). Using RT-PCR, in adult Ckm Cre; *Mbnl1*^fl/fl^, *Mbnl2*^fl/fl^ skeletal muscle tissue we found that only ∼5% of total *Limch1* mRNA included the six skeletal muscle-specific exons compared to ∼66% in age-matched WT skeletal muscle tissue (**Figure 6B, 6C**). Similarly, in adult Ckm Cre; *Mbnl1*^fl/fl^, *Mbnl2*^fl/fl^ skeletal muscle tissue we found significantly decreased mLIMCH1 protein expression and a reversion to the fetal predominant protein isoform, uLIMCH1 (**Figure 6D**). We conclude from the results that the MBNL family plays a significant role in the tissue-specific regulation of LIMCH1 and MBNL sequestration is the probable mechanism by which *LIMCH1* is mis-spliced in DM1.

**Figure 6.**
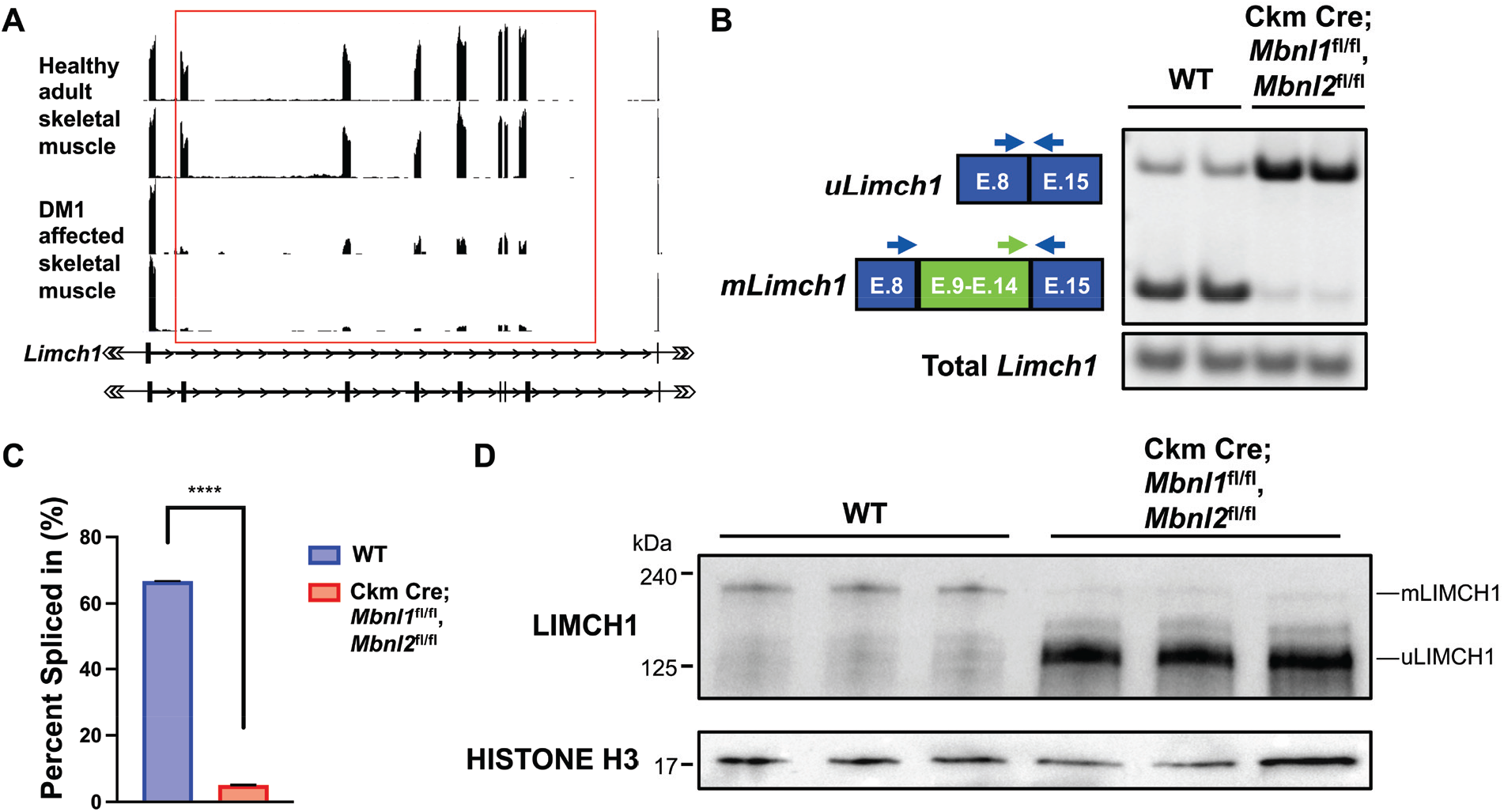
*LIMCH1* is mis-spliced in DM1 skeletal muscle. (A) RNA-seq tracks of *LIMCH1* from DM1 and non-DM1 human skeletal muscle. The red box indicates the muscle-specific region that is mis-spliced in DM1 skeletal muscle. (B) RT-PCR showing *Limch1* alternative splicing pattern in WT and Ckm Cre; *Mbnl1*^fl/fl^, *Mbnl2*^fl/fl^ skeletal muscle. (C) Quantification of *Limch1* muscle-specific exon inclusion in Ckm Cre; *Mbnl1*^fl/fl^, *Mbnl2*^fl/fl^ skeletal muscle (n= 2 WT, 2 Ckm Cre; *Mbnl1*^fl/fl^, *Mbnl2*^fl/fl^). (D) LIMCH1 protein expression in Ckm Cre; *Mbnl1*^fl/fl^, *Mbnl2*^fl/fl^ skeletal muscle (n= 3 WT, 3 Ckm Cre; *Mbnl1*^fl/fl^, *Mbnl2*^fl/fl^). Data represents the mean ± SEM and were analyzed using a two-tailed *t* test. ****P<.0001. DM1, myotonic dystrophy type 1; E, exon; WT, wild-type.

## Discussion

Alternative splicing is a mode of post-transcriptional gene regulation that contributes to the production of novel isoforms in a wide variety of tissues, thereby leading to the functional diversification of the proteome^36^. Further exploration is needed to understand the role of tissue-specific alternative splicing transitions and the pathological consequences of their mis-regulation in striated muscle. By screening for evolutionarily conserved and developmentally regulated alternative splicing transitions in skeletal muscle we identified a postnatal splicing transition in the *Limch1* gene. This unique splicing event is characterized by the coordinated inclusion of six exons constituting 454 in-frame amino acids and leads to the production of a novel skeletal muscle-specific isoform, mLIMCH1.

In this work, we sought to establish the contribution of mLIMCH1 to skeletal muscle function. We generated mice homozygous for a genomic deletion of the six exons encoding the muscle-specific isoform thereby exclusively expressing the fetal predominant isoform, uLIMCH1 in adult skeletal muscle. These mice demonstrated significant skeletal muscle weakness *in vivo* and deficits in skeletal muscle force generation *ex vivo*. We found decreased force generation in both slow-twitch muscle (soleus) and fast-twitch muscle (EDL).

Importantly, we saw no differences in muscle architecture based on histology and no muscle wasting based on muscle weights of the HOM *Limch1* 6exKO mice. These results provide evidence that global disruptions in muscle morphology are not contributing to the reduced force production and muscle weakness. It is likely that mLIMCH1 is contributing to normal muscle function through the maintenance of interfibrillar properties intrinsic to the muscle fiber.

To understand the specialized function of mLIMCH1, we used immunofluorescent microscopy to establish the isoform-specific endogenous localization of mLIMCH1 in the muscle myofiber. We found that in WT myofibers LIMCH1 had a preferential localization to the sarcolemma region with repeated staining observed at two micron intervals. To differentiate localization between mLIMCH1 and uLIMCH1 isoforms, we synthesized an mLIMCH1 isoform-specific antibody and found that mLIMCH1 is observed predominantly in the sarcolemma region. This localization is disrupted in HOM *Limch1* 6exKO myofibers where uLIMCH1 exhibited diffuse distribution throughout the myofiber underscoring the necessity of the skeletal muscle-specific isoform for normal LIMCH1 localization.

The disruption of calcium handling can alter skeletal muscle contraction in the absence of overt morphological changes^29^. The decreased rate of relaxation and rate of contraction in EDL from HOM *Limch1* 6exKO mice suggests calcium handling as a mechanism of the observed muscle weakness. With the gross morphology of skeletal muscle intact in HOM *Limch1* 6exKO, we show that mLIMCH1 is functionally significant for calcium handling with deficits in peak calcium release during myofiber stimulation likely involved in the resulting muscle weakness. T-tubule related proteins in the HOM *Limch1* 6exKO myofibers appear structurally intact, which necessitates future studies on the mechanism by which mLIMCH1 removal affects homeostatic calcium handling.

We also identified *LIMCH1* mis-splicing in DM1 muscle and showed that loss of MBNL activity, as observed in DM1, leads to the mis-regulation of the *Limch1* muscle-specific exons. Many genes are mis-spliced in DM1 and while some pathogenic splice variants are directly implicated in disease pathology, the contribution of other variants is not well understood. In this study, we did not directly examine the role *LIMCH1* is playing in DM1 progression. However, future studies will determine the degree to which *LIMCH1* mis-splicing contributes to strength and force loss in DM1.

There are certain factors from our study that should be taken into consideration. In HOM *Limch1* 6exKO skeletal muscle tissue, higher uLIMCH1 expression itself may contribute to skeletal muscle deficits since the same amount of pre-mRNA is produced and thus more uLIMCH1 protein will be produced than normally observed in adult tissue. Further, there is an inherent challenge of studying the protein binding characteristics of mLIMCH1 and structural contributions of mLIMCH1 to muscle cell architecture due to the lack of mLIMCH1 expression in cultured myoblasts or differentiated myotubes likely attributed to the immature state of these cells even when fully differentiated. With this study establishing the functional significance of mLIMCH1 in skeletal muscle, future work is focused on identifying the specific protein interactors of mLIMCH1 and uLIMCH1 facilitating skeletal muscle function both during development and in adult tissue.

## Materials and Methods

### Animals

For all animal studies, NIH guidelines for use and care of laboratory animals approved by Baylor College of Medicine Institutional Animal Care and Use Committee were followed. Heterozygous *Limch1* 6exKO mice were crossed to produce litters with homozygous *Limch1* 6exKO mice and WT mice. The WT mice are offspring from the founder line that established the *Limch1* 6exKO mouse line and were used in all of the experiments. All mice used in the experiments were 10-12 weeks old except when otherwise noted. Ckm Cre; *Mbnl1*^fl/fl^, *Mbnl2*^fl/fl^ tissue used in RT-PCR and western blot experiments were collected from a previously unpublished mouse line. In collaboration with the BCM Mouse Embryonic Stem Cell Core (mES) Core, loxP sites flanking exon three in *Mbnl1* and exon two in *Mbnl2* were introduced, and mice were appropriately crossed to produce *Mbnl1*^fl/fl^, *Mbnl2*^fl/fl^ mice. These mice were subsequently crossed with a Ckm Cre line (B6.FVB(129S4)-Tg(Ckmm-cre)5Khn/J; Jackson Labs 006475) to induce a knockout of Mbnl1 and *Mbnl2* during developmental expression of Ckm.

### CRISPR-Mediated *mLimch1* Deletion

*Limch1* 6exKO mice were developed in an FVB background in collaboration with the BCM mES Core. To delete the alternatively spliced exons 9-14 of *Limch1*, 4 single-guide RNAs (sgRNAs) were selected by the mES Core using the Wellcome Trust Sanger Institute Genome Editing website (http://www.sanger.ac.uk/htgt/wge/) positioned to flank the genomic region containing alternative exons 9-14 of *Limch1* (5′ sgRNA: http://www.sanger.ac.uk/htgt/wge/crispr/485343919, http://www.sanger.ac.uk/htgt/wge/crispr/485343920 and 3′ sgRNA: http://www.sanger.ac.uk/htgt/wge/crispr/485345259, http://www.sanger.ac.uk/htgt/wge/crispr/485345265).

Briefly, fertilized oocytes were collected and microinjected with the sgRNA/CRISPR-Cas9 mixture in ∼100 pronuclear stage zygotes. The injected zygotes were transferred into pseudopregnant females, at least 25 zygotes per recipient female. F0 mice were genotyped by standard PCR targeting the flanking region outside the six exons to confirm the null allele. F1 animals from two F0 founders were used to start two independent lines. The junctions of the genomic deletions in both lines were determined by sequencing PCR products generated across the junctions. Both lines were backcrossed to wild type FVB at least seven generations to out-cross off-target loci prior to performing experiments. Both lines showed comparable grip strength deficits and one line was selected experimentation.

### RT-qPCR and RNA splicing

Quadriceps skeletal muscle tissues were dissected, flash frozen in liquid nitrogen, and maintained at -80° C prior to RNA isolation. Total RNA was isolated from skeletal muscle tissues using TRIzol reagent (catalog number 15596018, Invitrogen) and cDNA was prepared from 1 μg of DNase treated RNA (catalog number 4368813, Thermo Fisher) followed by PCR amplification. For analysis of alternative splicing events, primers annealing to flanking constitutive exons were designed as follows: To amplify *uLimch1* and *mLimch1*-F1: 5’-AGCGGGAGGAATTCAGGAAG-3’, R1: 5’-CGTCAATTCCCTCCTCCTCT-3’, R2: 5’-TCCTCACACCGCATGTCAAA-3’. To amplify total *Limch1*-F1: 5’-GAAGACGGTGAAGAAAGGCC-3’, R2: 5’-GTTGTCGTAAAGTGCTGGGG-3’. PCR products were resolved on a 5% polyacrylamide gel to determine both the total *Limch1* RNA and the percentage of *uLimch1* and *mLimch1* RNA. Ethidium bromide stained RT-PCR bands were analyzed using Kodak Gel Logic 2000 and Carestream software. Percent Spliced In (PSI) was calculated to determine the percent of mRNA from the gene that contains the alternative exons, quantified using densitometry according to the equation: PSI = 100 X [Inclusion band/(Inclusion band + Skipping band)].

### Protein isolation and immunoblotting

Tissue protein extracts were prepared from isolated skeletal muscle tissue by mechanical homogenization with a bullet blender (Next Advance) in 1X RIPA lysis buffer (catalog number 9806, Cell Signaling) supplemented with 1X Xpert Protease Inhibitor Cocktail Solution (catalog number P3100, GenDEPOT) with subsequent centrifugation to remove cell debris (15,000*g* for 15 minutes at 4°C). BCA protein assay kit (catalog number PI23225, Thermo Fisher) was used to quantify protein concentration and protein samples were diluted to 3 μg/μL in Laemmli SDS sample buffer (catalog number J61337.AC, Thermo Fisher) and boiled at 100°C for 5 min. 15-30 μg of protein lysate was loaded and separated on 12% Tris-glycine SDS-PAGE gels and transferred to nitrocellulose membranes (catalog number 1620167, Bio-Rad) using Trans-Blot Turbo Transfer System (Bio-Rad) for western blot analysis. Membranes were blocked with 5% milk in PSB-T (0.1% Tween-20, Sigma) for one hour and incubated overnight at 4°C in 5% milk/PBS-T with the following antibodies: anti-LIMCH1 (1:400, catalog number HPA063840, Sigma), anti-MBNL1 (1:1,000, catalog number 3A4: sc-47740, Santa Cruz Biotechnology), anti-MBNL2 (1:1,000, catalog number 3B4: sc-136167, Santa Cruz Biotechnology), anti-GAPDH (1:10,000, catalog number 14C10: 2118, Cell Signaling), anti-Histone H3 (1: 2,000, catalog number D1H2: 4499, Cell Signaling), and anti-mLIMCH1 (1:400, Biomatik). Membranes were washed three times with PBS-T and incubated for one hour at room temperature in HRP-conjugated goat anti–rabbit or goat anti-mouse secondary antibody (1:10,000, Jackson Immunoresearch) in 5% milk/PBS-T after which membranes were washed three times with PBS-T. Immunoreactivity was detected using West-Q Pico Dura ECL Solution (Catalog number W3653, GenDEPOT) and membranes were imaged on a ChemiDoc XRS+ Imaging system (Bio-Rad).

### Grip strength and body weight

A grip strength meter (Columbus Instruments, Columbus, OH) was used to conduct an all-limb grip strength assessment on WT and *Limch1* 6exKO FVB mice at 10-12 weeks of age. Mice were acclimated in the testing room for at least 10 minutes prior to testing to minimize effects of stress on behavior during testing. Mice genotypes were blinded to the experimenter and the same experimenter conducted all grip strength assessments to reduce variability. As the mouse grasped the grid, the peak pull force in grams was recorded on a digital force transducer. The procedure was repeated for a total of three trials separated by 15 minute inter-trial intervals. At the end of trial three, animals were weighed and grip force was standardized to the respective body weight of the individual mouse.

### Skeletal muscle weights

Muscle weights of gastrocnemius, quadriceps, and soleus was measured on an analytical balance and normalized to tibia length at the conclusion of the grip strength assessment. Direct comparisons of normalized muscle weights were made between age-matched treatment groups.

### Histology

Skeletal muscle from quadriceps tissue was isolated, connective tissue removed, fixed in 10% formalin for 24 hours, paraffin-embedded and cut in 10 μm cross sections. Hematoxylin and eosin staining (H&E) and picrosirius staining of skeletal muscle was performed using standard procedures. Images were acquired using an Olympus BX41 microscope with Olympus DP70 camera.

### Isometric ex vivo force analysis

Soleus and EDL muscle were dissected from male mice and using silk suture (4-0), one tendon of the muscle was tied to a fixed hook and the other tendon of the muscle was tied a force transducer (F30, Harvard Apparatus). The muscle was maintained in a physiological saline solution and supplemented with 95% O_2_–5% CO_2_ at 30 °C. Optimal length of the muscle (Lo) was determined to produce maximum twitch force, while optimal voltage was determined following Lo. Muscles were allowed to equilibrate for 10 minutes following Lo and optimal voltage assessment. Pulse and train durations of 0.25 and 400 ms, respectively, were used to assess contractile properties of the muscle with current passing through two platinum electrodes (PanLab LE 12406, Harvard Apparatus). Kinetics, force-frequency, fatigue, and recovery from fatigue protocols were collected and analyzed in LabChart 8 (ADInstruements) with the Peak Analysis module. Force-frequency was tested by stimulating the muscle at 1 Hz, 5 Hz, 10 Hz, 20 Hz, 40 Hz, 60 Hz, 80 Hz, 120 Hz, 150 Hz and 200 Hz with one minute intervals between each frequency tested. After a two minute rest period, the fatigue protocol was conducted by stimulating the soleus with a 40 Hz stimulus and the EDL with a 70 Hz stimulus every two seconds for five minutes. After a five minute rest period, the recovery from fatigue protocol was assessed by stimulating the muscle with alternating frequencies of 40 Hz and 150 Hz every minute for 10 minutes. At the end of the recovery protocol, muscle length was measured using a hand-held electronic caliper, muscles were removed from the connected apparatus and excess connective tissue was removed and muscles were blotted dry and weighed. For force frequency, the muscle weight and Lo recorded were used to calculate absolute force expressed as (N/ cm^2^)^37^. Rate of contraction and rate of relaxation was collected during the force-frequency protocol during a single twitch (1 Hz) and was based on the slope of the force curve. The muscle weight and Lo recorded were used to determine the forces expressed as a function of time (N/ cm^2^/s) for rate of recovery (-dp/dt) and rate of relaxation (dp/dt). For the fatigue protocol, force was normalized to the first stimulus prior to fatigue. For the recovery protocol, force was normalized to a 40 Hz or 150 Hz stimulus prior to fatigue.

### FDB myofiber isolation

Mice were anesthetized with 2% isoflurane and euthanized with subsequent cervical dislocation. FDB tissue was surgically dissected and connective tissue was removed. The FDB muscle was placed into a 35 mm dish containing Minimum Essential Medium (catalog number 51411C, Sigma), 2 mg/mL collagenase type 1 (catalog number C0130, Sigma), 0.1% gentamyacin, and incubated in 5% CO2 at 37° C for two hours. After incubation, the FDB muscle was washed with PBS, transferred to a new, non-coated 35 mm dish and single fibers were gently released with serial trituration into 2 mL of DMEM (catalog number 11-965-092, Thermo Fisher) supplemented with 10% fetal bovine serum (FBS, catalog number F0900-050, GenDEPOT), 1% penicillin / streptomycin and incubated in 5% CO2 at 37° C overnight. The isolated myofibers were then used for subsequent assays.

### Cell culture and *in vitro* transfections

C2C12 cells were cultured in 96-well plates (catalog number 89626, Ibidi) pre-coated with ECM Gel (catalog number E1270, Sigma) and maintained in DMEM supplemented with 10% FBS, 1% penicillin / streptomycin and incubated in 5% CO2 at 37° C. Once C2C12 cells reached 70-80% confluency, cells were transfected with *uLimch1* and *mLimch1* epitope tagged expression plasmids using Lipofectamine 3000 following the manufacturer’s protocol (catalog number L3000015, Invitrogen). *mLimch1* and *uLimch1* epitope tagged expression plasmids were commercially synthesized by GenScript. A Myc tag was included in the C-terminus region of the *uLimch1* sequence, while a Myc tag was included in the C-terminus region of the *mLimch1* sequence in addition to a FLAG tag which was inserted within the muscle-specific sequence of *mLimch1* in exon 10. The *uLimch1* and *mLimch1* sequences were cloned into the backbone of a pcDNA3.1 plasmid by GenScript. Transfected cells were incubated in antibiotic free DMEM supplemented with 10% FBS for 48 hours. Cells were then fixed in 10% formalin for 10 minutes and prepared for immunofluorescence and imaging.

### mLIMCH1 antibody synthesis

The mLIMCH1-specific primary antibody was commercially generated by Biomatik. The following protein antigen sequence was submitted to Biomatik for antibody synthesis-LEQAGIKVMPAAQRFASQKQLSEEKEAIRDIVLRKENSFLTHQHGNDSEAEGEVVCRLPDLEKDDFAARRARMNQTKPMVPLNQLLYGPY. Rabbits underwent multiple immunizations with the antigen which was expressed as a recombinant protein, to yield an immunogenic response. After multiple immunizations, the serum titer reached at least 1:8,000, and 3-5 mg of antibody was purified with antigen affinity purification. The resulting antibody was validated by Biomatik with ELISA (>1:32,000) and validated for western blot and immunofluorescence by our lab.

### Immunofluorescence

96-well plates (Ibidi) were coated with ECM Gel for 30 minutes at room temperature. 100 uL of isolated myofibers were transferred into the coated wells and incubated in 5% CO2 at 37° C for one hour to allow for myofiber adhesion. Myofibers were washed with PBS and fixed with 4% paraformaldehyde in PBS for 10 minutes at room temperature. Fixed myofibers were washed with PBS three times and permeabilized using 0.1% triton-X 100 (Sigma) in PBS for 10 minutes. Myofibers were then blocked in 5% FBS, 0.1% triton-X 100 in PBS at room temperature for one hour and incubated overnight in 5% FBS and 0.1% triton-X 100 in PBS at 4°C with the following antibodies: anti-LIMCH1 (1:200, catalog number HPA063840, Sigma), anti-mLIMCH1 (1:200, Biomatik), anti-RYR1 (1:200, catalog number 34C, Developmental Studies Hybridoma Bank), anti-BIN1 (1:200, catalog number 14647-1-AP, Proteintech). Transfected C2C12 cells were fixed and stained following the same protocol and incubated overnight in 5% FBS and 0.1% triton-X 100 in PBS at 4°C with the following antibodies: anti-mLIMCH1 (1:200, Biomatik), anti-Myc (1:200, 71D10: 2278, Cell Signaling), anti-DDDDK (1:200, catalog number ab1257, abcam). The following day, the myofibers or cells were washed three times with PBS and incubated in goat anti-rabbit 555 Alexa Fluor-conjugated secondary antibody (1:1000, catalog number A27039, Thermo Fisher) or goat anti-mouse 488 Alexa Fluor-conjugated secondary antibody (1:1000, catalog number A11029, Thermo Fisher) for two hours in 5% FBS and 0.1% triton-X 100, and washed three times with PBS. Transfected C2C12 cells were stained with DAPI (0.5 μg/mL, catalog number D9542, Sigma) for five minutes followed by one wash with PBS before imaging. For T-tubule staining with FM 4-64 (Catalog number T13320, Invitrogen) unfixed myofibers were loaded with 5 μg/mL of FM 4-64 dye and imaged immediately.

### Microscopy

Confocal microscopy on myofibers was performed using a Nikon A1-Rs inverted laser scanning microscope with a 60x Plan Apo VC / 1.4 NA oil-immersion objective. Transfected C2C12 cells images were acquired using a DeltaVision Elite (GE Healthcare) and processed using SoftWoRx software (GE Healthcare).

### Image processing and quantitative analysis

Plot profiles and image analytics were performed using Fiji (ImageJ). A mask was generated delimiting each myofiber by applying a threshold manually. The identified signal was then projected onto the original image file outlining the area for analysis. In the case of LIMCH1 immunofluorescence, the standard deviation of the immunofluorescence signal was determined as a means to measure the signal distribution of LIMCH1 throughout the entire myofiber of a single image plane. Two horizontal slices were chosen near the center of the myofiber and the standard deviation of immunofluorescence from each slice was averaged for each sample analyzed. Multiple myofibers from one mouse was analyzed and averaged to serve as a single sample with 6 mice (n=6) used for both the WT and *Limch1* 6exKO groups. Representative plot profiles were recorded from HOM *Limch1* 6exKO and WT myofibers through the myofiber (sarcolemma to sarcolemma) and along the sarcolemma in the center plane of the myofiber.

### Cellular Ca^2+^ imaging

FDB muscle was surgically dissected and individual myofibers were isolated as described above. 22×22 mm micro cover glass (catalog number 48366227, VWR) was coated with ECM Gel for 30 minutes at room temperature. 100 uL of isolated myofibers were transferred onto the coated cover glass slides and incubated in 5% CO2 at 37° C for 30 minutes to allow for myofiber adhesion. Once myofibers adhered, physiological saline solution (1.8 mM CaCl2, 120 mM NaCl, 4.7 mM KCl, 0.6 mM MgSO4, 1.6 mM NaHCO3, 0.13 mM KH2PO4, 7.8 mM glucose, 20 mM HEPES, pH 7.3) was added in a buffer exchange and 5 μM Fura-4F AM (catalog number F14175, Invitrogen) was added to the myofibers for 30 minutes at room temperature. Myofibers were washed with physiological saline solution at least 3 times to remove Fura-4F AM and allowed to de-esterify for 30 minutes. To determine optimal calcium response, the focal plane was adjusted and calcium transients were measured following twitch (1 Hz), 20 Hz tetanus, and 100 Hz tetanus with pulse and train durations of 0.5 and 250 ms, respectively. IonOptix Myocyte Calcium and Contractility Recording System (IonOptix, Westwood, MA) was used to monitor Fura-4F excitation (360/380 nm) and emission (510 nm). Intracellular calcium changes during myofiber stimulation were inferred from the recorded 360 nm / 380 nm ratio signals. Multiple myofibers from one mouse was analyzed and averaged to serve as a single sample with each sample (n) derived from different mice.

### Statistics

All quantitative experiments have at least three independent biological replicates with the exception of the RT-PCR of Ckm Cre; *Mbnl1*^fl/fl^, *Mbnl2*^fl/fl^ and WT skeletal muscle. Results are presented as mean ± standard error of the mean (SEM). All statistical analyses were calculated using Prism software (version 8.0, GraphPad) and methods and samples sizes used are shown in respective figure legends. p values were calculated by a two-tailed *t* test for all the experiments and a *P* value of less than 0.05 was considered statistically significant.

### Study approval

All mouse experiments were carried out in accordance with the *Guide for the Care and Use of Laboratory Animals* (National Academies Press, 2011) and approved by the Baylor College of Medicine Institutional Animal Care and Use Committee.

## Acknowledgements

Imaging for this project was supported by the Integrated Microscopy Core at Baylor College of Medicine and the Center for Advanced Microscopy and Image Informatics (CAMII) with funding from NIH (DK56338, CA125123, ES030285), and CPRIT (RP150578, RP170719) with a special thanks to Hannah Johnson and Fabio Stossi for their assistance and guidance. This project was supported by the Mouse Metabolism and Phenotyping Core at Baylor College of Medicine which is subsidized as a member of the institution’s Advanced Technology Cores, and through funding from the NIH (UM1HG006348, R01DK114356, R01HL130249). This project was supported by the Genetically Engineered Rodent Model (GERM) Core at BCM. The GERM Core is funded in part by the National Institutes of Health Cancer Center Grant (P30 CA125123). We also thank Ashish Rao for his valuable input on experimental design and manuscript review. We thank Riya Thomas and Jennifer Graham for their help with the grip strength assessment. We also thank members of the Cooper lab for their valuable discussions and help throughout the project. Images were created with BioRender. This project was supported by NIH grants R01 AR060733 (T.A.C.), R01 HL147020 (T.A.C.), F31AR078646 (M.S.P), RO1 AR061370 (G.G.R.), and by generous support from the Baylor Research Advocates for Student Scientists (BRASS) who provided financial support (M.S.P) for many of the experiments conducted in this study.

## Author contributions

M.S.P and T.A.C. developed the project. M.S.P, G.G.R., and T.A.C. designed the experiments. R.H. conducted RT-PCR experiments, M.S.P performed the rest of the experiments with valuable input from T.A.C. and G.G.R. All authors contributed to the interpretation of results and production of the final manuscript.

---

**Figure S1.**
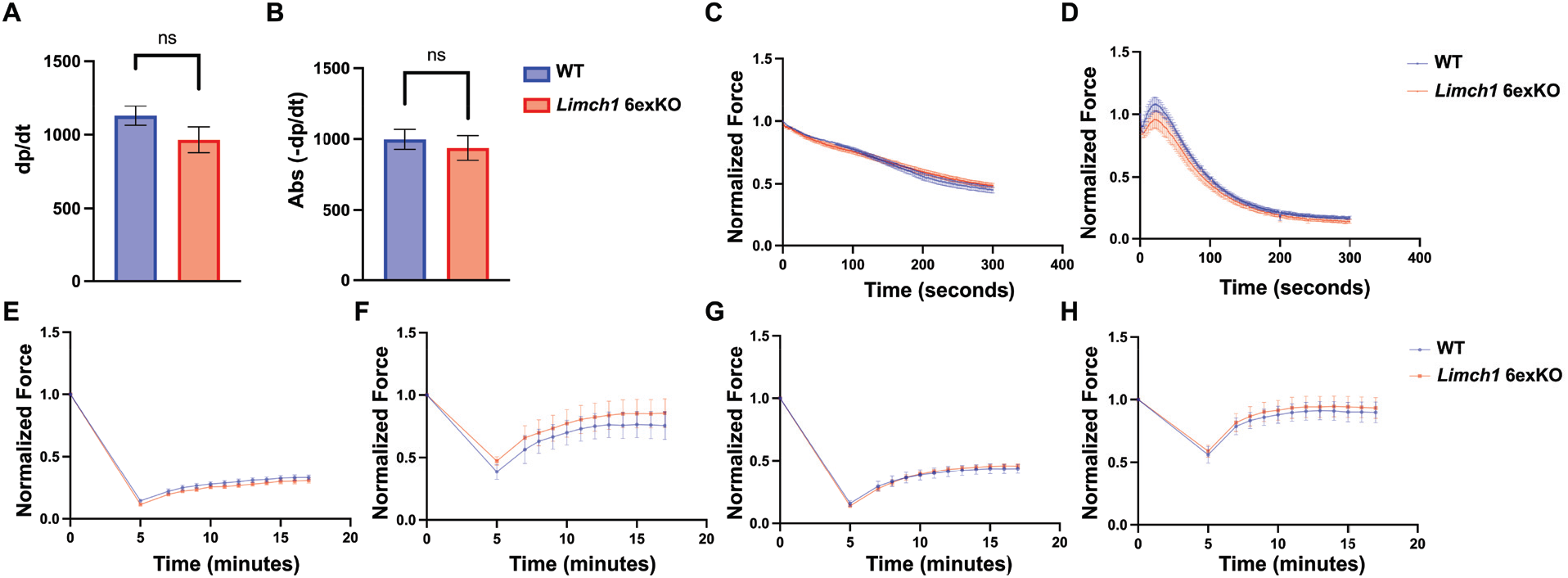
Fatigue and recovery during *ex vivo* isometric contraction remain unchanged in HOM *Limch1* 6exKO skeletal muscle. (A) Rate of contraction (dp/dt) in the soleus following an *ex vivo* isometric twitch. (B) Rate of relaxation (-dp/dt) in the soleus following an *ex vivo* isometric twitch. (C) Normalized force curve after stimulating the soleus to induce fatigue. (D) Normalized force curve after stimulating the EDL to induce fatigue. (E) Recovery of low frequency (40 Hz) force in the EDL. (F) Recovery of low frequency (40 Hz) force in the soleus. (G) Recovery of high frequency (150 Hz) force in the EDL. (H) Recovery of high frequency (150 Hz) force in the soleus. (EDL: n= 6 WT, 8 HOM; soleus: n= 6 WT, 6 HOM). Data represents the mean ± SEM and were analyzed using a two-tailed *t* test for each time point assessed. WT, wild-type; EDL, extensor digitorum longus; HOM, homozygous.

**Figure S2.**
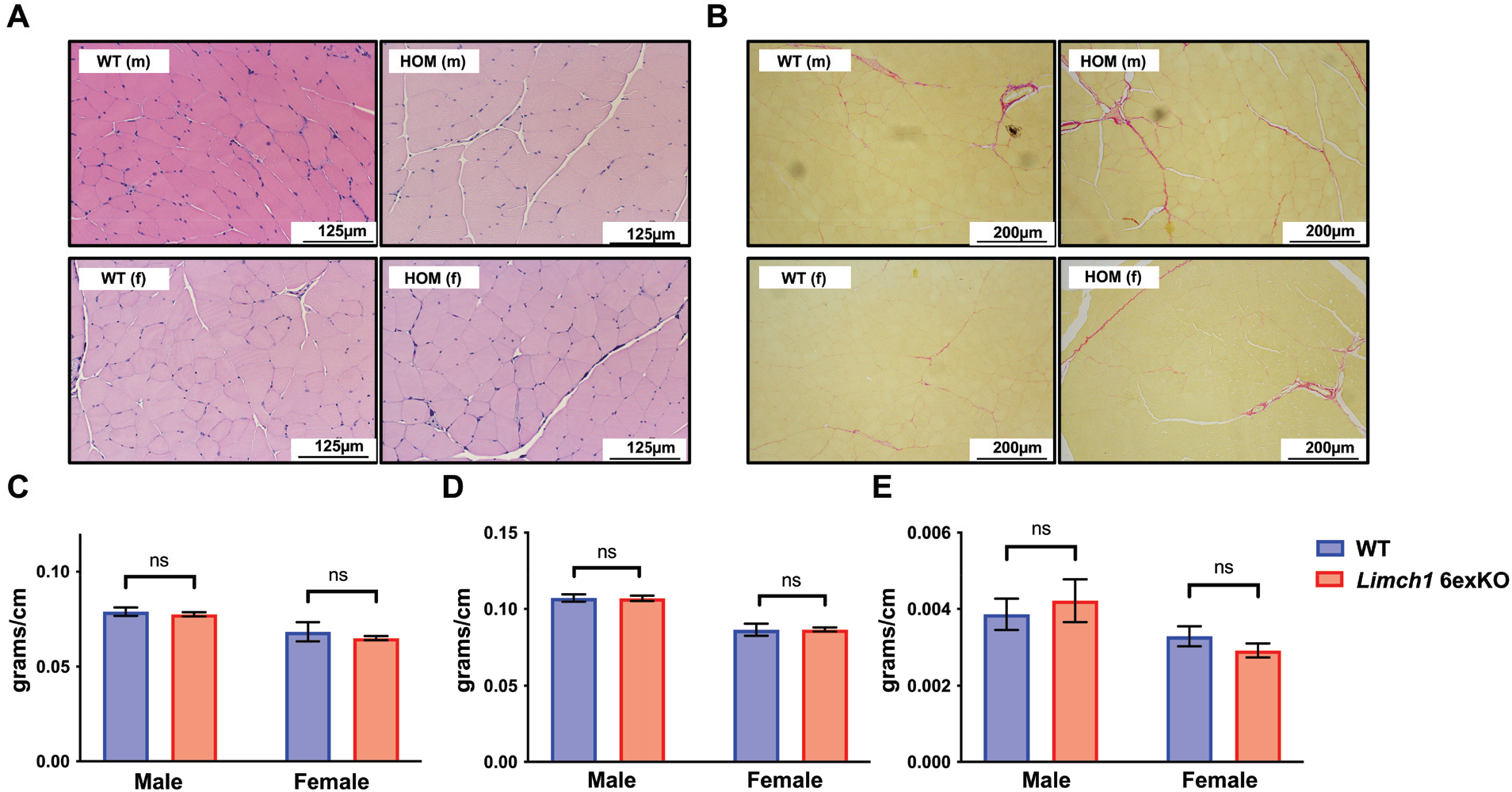
Gross morphological changes are not observed in HOM *Limch1* 6exKO skeletal muscle tissue. (A) Skeletal muscle cross sections stained with H&E of male and female HOM *Limch1* 6exKO and age-matched WT tissue. (B) HOM *Limch1* 6exKO and age-matched WT skeletal muscle cross sections stained with picrosirius stain. (C-E) Muscle weights divided by tibia length from WT and HOM *Limch1* 6exKO tissue. (C) Gastrocnemius (D) Quadriceps (E) Soleus (n= 13 WT, 11 HOM males; n= 11 WT, 20 HOM females). (ns=not significant). Data represents the mean ± SEM and were analyzed using a two-tailed *t* test. HOM, homozygous; m, male; f, female; WT, wild-type; ns, not significant.

**Figure S3.**
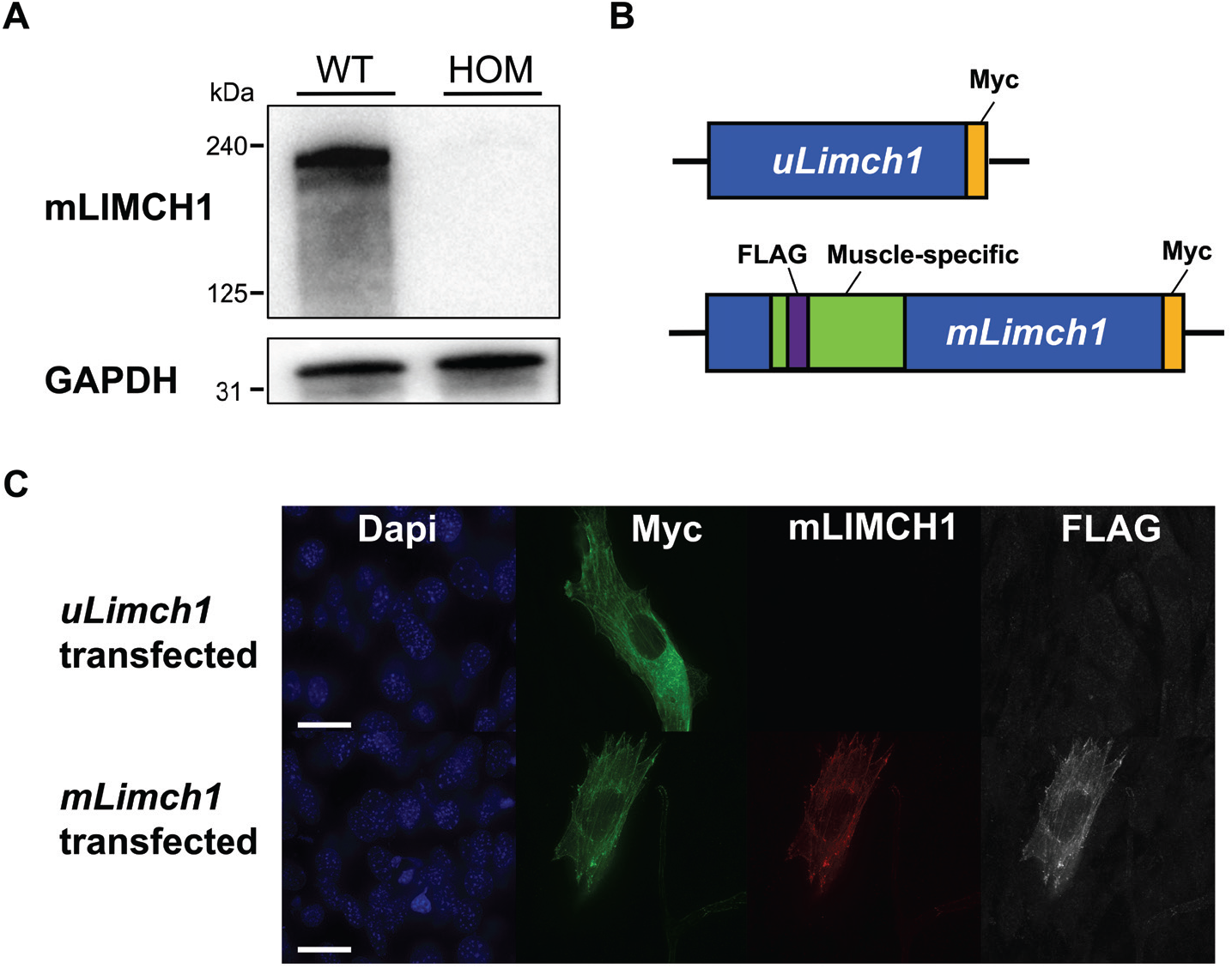
mLIMCH1-specific antibody recognizes mLIMCH1 in western blot and immunofluorescence assays. (A) Western blot of adult HOM *Limch1* 6exKO and WT skeletal muscle probed with mLIMCH1-specific antibody. **(B)** Graphic illustration of *uLimch1* and *mLimch1* epitope-tagged expression plasmids with a myc epitope tag at the C-terminus and a FLAG epitope tag inserted within exon 10 of the muscle-specific region of *mLimch1*. (C) Immunofluorescence staining of uLIMCH1 and mLIMCH1 transfected C2C12 myoblasts. Scale bar, 15 μM. HOM, homozygous; WT, wild-type.

**Figure S4.**
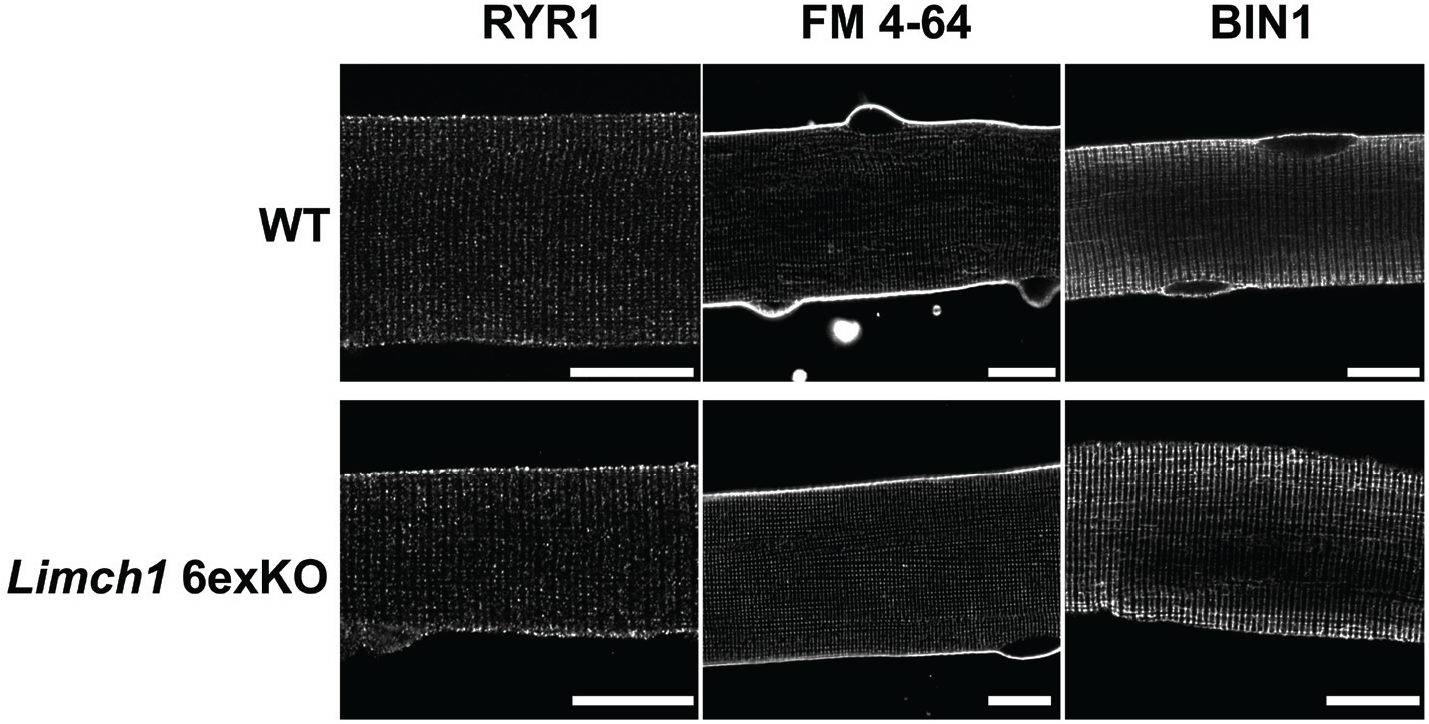
T-tubule structure remains intact in HOM *Limch1* 6exKO myofibers. Immunofluorescence staining of RYR1, FM 4-64 (T-tubule membrane), and BIN1 in *Limch1* 6exKO and WT age-matched control myofibers. Scale bar, 15 μM. WT, wild-type; HOM, homozygous; T-tubule, transverse tubule.

**Figure S5.**
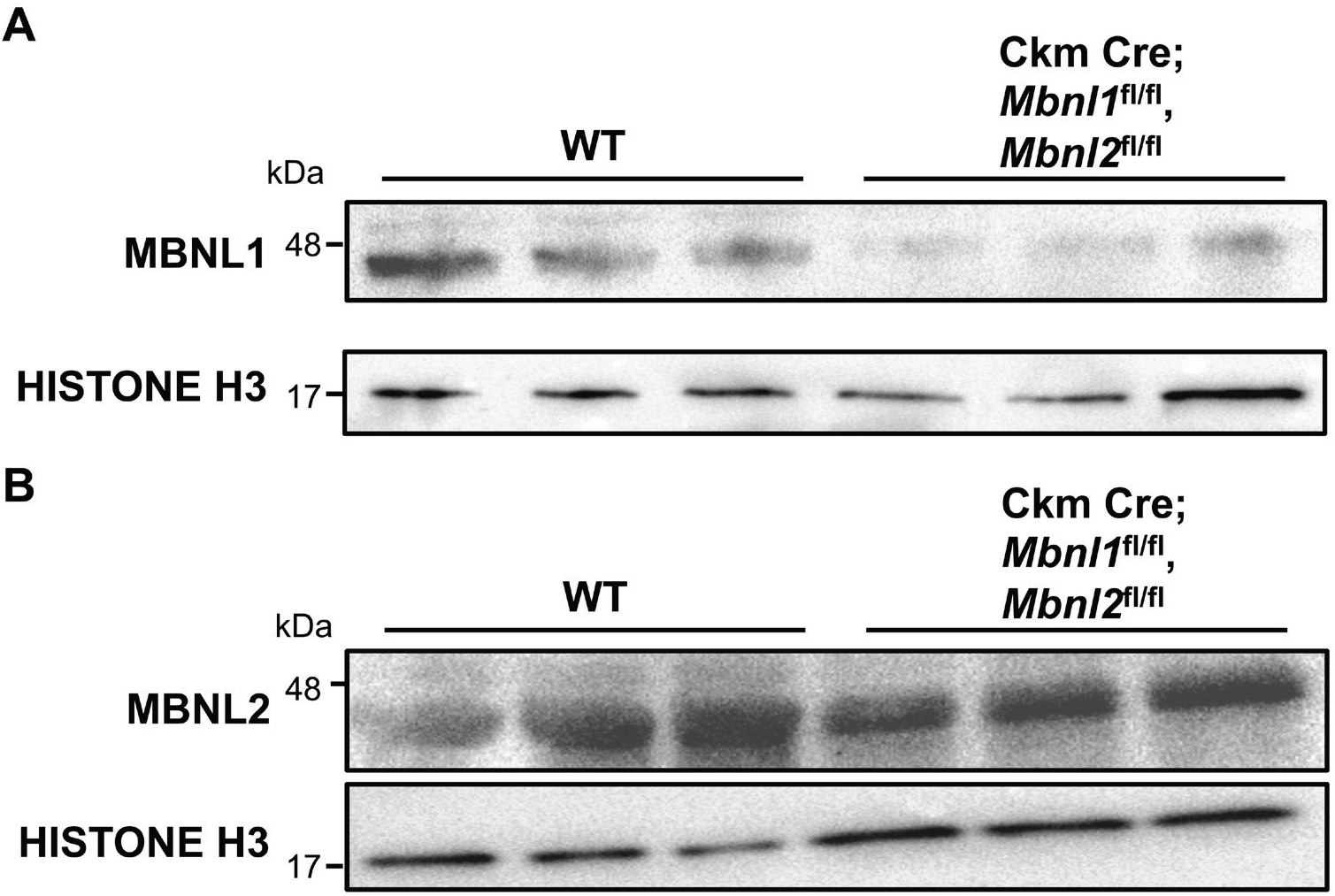
Ckm Cre; *Mbnl1*^fl/fl^, *Mbnl2*^fl/fl^ skeletal muscle shows decreased levels of MBNL1 and MBNL2. (A) MBNL1 protein expression in Ckm Cre; *Mbnl1*^fl/fl^, *Mbnl2*^fl/fl^ skeletal muscle (n= 2 WT, 2 Ckm Cre; *Mbnl1*^fl/fl^, *Mbnl2*^fl/fl^). (B) MBNL2 protein expression in Ckm Cre; *Mbnl1*^fl/fl^, *Mbnl2*^fl/fl^ skeletal muscle. WT, wild-type (n= 2 WT, 2 Ckm Cre; *Mbnl1*^fl/fl^, *Mbnl2*^fl/fl^).

